# Galectin-8 regulates primary cilium in hypothalamic neurons through an L-type calcium channel/Aurora kinase A/HDAC6 pathway impacting body energy balance

**DOI:** 10.64898/2026.04.09.716665

**Authors:** Cristian Herrera-Cid, María Paz Hernández, Daniela Pinto, Alejandra Aranguiz, Francisca Pérez-Molina, Ariel Vivero, Daniela Cortés-Díaz, Claudia Jara, Sofia Espinoza, Andrea Soza, Cheril Tapia-Rojas, Bredford Kerr, Eugenia Morselli, Alfonso González

**Affiliations:** Centro de Biología Celular y Biomedicina (CEBICEM) Facultad de Medicina y Ciencia, Universidad San Sebastián, Santiago, Chile; Departamento de Ciencias Básicas, Facultad de Medicina y Ciencia, Universidad San Sebastián, Santiago, Chile; Centro Científico y Tecnológico de Excelencia (CCTE) Ciencia & Vida

**Author notes:** Corresponding author: Alfonso González, M.D., Ph.D., Centro de Biología Celular y Biomedicina, Facultad de Medicina y Ciencia, Universidad San Sebastián, Santiago, Chile., Address: Lota 2465, Santiago 7510157, Chile.

## Abstract

**OBJETIVE:** Food intake, energy expenditure, and metabolic homeostasis depend on hypothalamic neurons’ responses to peripheral signals, such as leptin, involving the primary cilium (PC). The PC is crucial for signal transduction and is dynamically regulated by assembly/disassembly or reabsorption of its microtubules-based axoneme. Absence or reduction in the length of PC is associated with obesity and type-2 diabetes (T2D). In other cellular systems, PC reabsorption is primarily regulated by calcium-mediated activation of the Aurora kinase A (AurkA)/histone deacetylase C6 (HDAC6) axis, which promotes axonemal disassembly. Here, we explore the role of Galectin-8 (Gal-8), a glycan-binding protein, in regulating PC structure and signaling related to metabolic parameters in hypothalamic neurons.

**METHODS:** Gal-8 effects were assessed in hypothalamic Clu-177 cells by analyzing the PC presence and length by immunofluorescence, PC dynamics, and intracellular calcium changes by *in vivo* cell imaging, activation of FAK, Src, AurkA, HDAC6 and STAT3 by immunoblot, and Gal-8 interactions with β1-integrins by pull-down assays. Gal-8-KO mice were used to evaluate PC length in hypothalamic neurons, metabolic phenotype, and responses to Gal-8 intranasal administration.

**RESULTS:** In Clu-177 cells, Gal-8 induced PC reabsorption and reduced responsiveness to leptin signaling towards STAT3 activation. PC reabsorption involves glycan-mediated Gal-8 interactions with a5b1 and a3b1 integrins, activation of FAK and Src leading to calcium influx through L-type calcium channels (LTCC), and subsequent AurkA/HDAC6 axis activation. Gal-8-KO mice showed longer PC in hypothalamic neurons, higher STAT3 activation, decreased body weight and food intake, improved glucose tolerance, higher locomotor activity, and a glycolytic respiratory exchange rate (RER). Daily intranasal Gal-8 administration for 4 days restored hypothalamic PC length and STAT3 signaling, as well as RER in Gal-8-KO mice to the level of WT mice.

**CONCLUSIONS:** Endogenous Gal-8 is required to maintain PC structure and leptin signaling in hypothalamic neurons, impacting body weight, energy balance, and glucose homeostasis. The mechanism involves calcium influx via LTCC downstream of b1-integrin/FAK/Src signaling and subsequent AurkA/HDAC6 axis activation. Both Gal-8 and the AurkA/HDAC6 axis may offer new therapeutic opportunities for treating metabolic diseases characterized by ciliogenesis impairment, including obesity and type-2 diabetes.

## INTRODUCTION

The hypothalamus regulates energy balance by integrating peripheral signals with the central functions of distinct neuronal populations that control appetite and maintain body weight (1, 2). One of the most extensively studied systems is based on discrete groups of neurons concentrated in the arcuate nucleus (ARC) of the hypothalamus and their elaborated circuitries that drive anorexigenic responses to leptin, a hormone secreted by white adipose tissue, particularly after food intake, to induce satiety (3–6). ARC neurons express the long isoform of the leptin receptor, LepRb, which signals through multiple pathways, including tyrosine phosphorylation and activation of STAT3 (7). Alterations in this system due to impaired expression or functional deficiency of leptin or LepRb, as well as various forms of leptin resistance, have long been associated with obesity (7–10). Obesity represents a major public health problem because of its pandemic prevalence worldwide and its predisposition to develop other diseases, such as cardiovascular disorders and type 2 diabetes (11, 12). However, the mechanisms that determine the magnitude of the neural responses to leptin and their dysfunctions that lead to or are associated with obesity have not been fully elucidated (1, 7). It is therefore important to identify endogenous factors in the brain that may affect the sensitivity of hypothalamic ARC neurons to leptin. One aspect that has not been sufficiently explored in this context is the regulation of primary cilium structure, whose features influence the strength of leptin-evoked cellular signaling (13, 14).

The primary cilium (PC) is an antenna-like organelle that protrudes from the surface of most mammalian cells and functions as a sensor of extracellular signals (15). Structurally, it consists of a specialized membrane contiguous with the plasma membrane and a microtubule-based axoneme that extends from the mother centriole of the basal body (15). The PC is highly dynamic, undergoing changes in length or complete loss through tightly regulated processes of assembly, disassembly, and shedding in response to extracellular cues (15–18). This dynamic behavior provides an additional layer of control over intracellular signaling pathways. In neurons, alterations in PC structure have been linked to changes in signaling strength, including pathways mediated by receptors such as those for leptin and insulin (14, 19, 20). Importantly, defects in PC structure or function underlie a group of disorders known as ciliopathies, which are frequently associated with hyperphagia, early-onset obesity, and type 2 diabetes (13).

Consistent with these observations, the length of the PC in hypothalamic neurons is variable and influenced by dietary conditions and obesity-related states (13, 14, 19), suggesting that endogenous factors regulate the balance between ciliogenesis and PC resorption. Among the signaling pathways affected by PC dynamics, leptin signaling plays a central role in the regulation of energy balance. The hypothalamus integrates peripheral metabolic cues with central neuronal circuits to control appetite and body weight (1, 2). Within this system, neurons located in the arcuate nucleus (ARC) respond to leptin—a hormone secreted by white adipose tissue—by activating anorexigenic pathways that promote satiety (3–5). These neurons express the long isoform of the leptin receptor (LepRb), which signals through multiple pathways, including tyrosine phosphorylation and STAT3 activation (7). Disruptions in leptin production, LepRb function, or downstream signaling are strongly associated with obesity and related metabolic disorders (7–10), a major global health concern due to its increasing prevalence and association with cardiovascular disease and type 2 diabetes (11, 12). Notably, the structural state of the PC has been shown to influence leptin signaling strength in ARC neurons (13, 14). While leptin itself promotes ciliogenesis (14, 20), ciliary dysfunction can lead to leptin resistance even before the onset of obesity (21). However, the endogenous mechanisms that regulate PC maintenance, remodeling, and resorption in these neurons—and how these processes affect leptin responsiveness—remain poorly understood.

Galectin-8 (Gal-8) is a 34 kDa carbohydrate-binding protein broadly expressed across tissues (22–24), including multiple regions of the brain such as the hypothalamus (25–27). Members of the galectin family share conserved carbohydrate-recognition domains (CRDs) with affinity for β-galactosides and are secreted through non-classical pathways (23, 25). Gal-8 contains two distinct CRDs: an N-terminal domain with preferential binding to α2-3-sialylated glycans and a C-terminal domain that recognizes non-sialylated oligosaccharides, connected by a linker peptide that defines different isoforms (22, 28, 29). Through interactions with cell-surface glycoproteins, Gal-8 regulates diverse cellular processes (22, 23, 25, 30). Among its best-characterized binding partners are β1-integrins, particularly α5β1 and α3β1, whose activation has been shown to increase intracellular calcium levels (23, 27, 31–37). Independently, increases in cytosolic calcium have been linked to PC shortening via activation of calmodulin and Aurora A kinase (AURKA) signaling in both mitotic and non-mitotic cells (38–42). Whether this pathway operates in neurons, however, remains unknown.

In the brain, Gal-8 is expressed in multiple regions, including the hypothalamus (26, 27). In addition to local expression, Gal-8 may also reach the hypothalamus by diffusion from adjacent regions such as the thalamus, as well as from the cerebrospinal fluid (CSF) produced by the choroid plexus, two sites that display the highest levels of Gal-8 expression (26, 27). Components of the CSF can access most regions of the cerebral parenchyma via direct transcytosis across permeable ependymal cells and/or through the glymphatic cerebrospinal fluid–interstitial fluid (CSF–ISF) circulation system (43). Variable levels of Gal-8 have been detected in human cerebrospinal fluid (26). However, to date, the only functions reported for Gal-8 in the brain are neuroprotective and immunosuppressive roles (26, 27). Whether Gal-8 influences hypothalamic neuronal signaling and energy balance regulation has not been explored.

Here, we provide evidence that Gal-8 is a key regulator of primary cilium dynamics and energy balance. Using Gal-8 knockout (Gal-8 KO) mice, we show that endogenous Gal-8 is required to maintain body weight, adipose tissue mass, glucose homeostasis, and the structural integrity of the PC in ARC neurons. Gal-8 deficiency leads to increased PC length and enhanced hypothalamic STAT3 phosphorylation, both of which are reversible upon intranasal Gal-8 administration. Mechanistically, using Clu-177 cells as a model of adult hypothalamic neurons (44), we demonstrate that Gal-8 promotes PC resorption through interaction with β1-integrins, triggering calcium influx and activation of AURKA. This ciliary remodeling is associated with reduced leptin responsiveness. Together, these findings identify Gal-8 as a previously unrecognized regulator of ciliogenesis in hypothalamic neurons, linking extracellular glycan recognition to the control of energy balance.

## RESULTS

### Primary cilium length in ARC neurons is increased in Gal-8 KO mice and normalized by intranasal Gal-8 administration

To assess whether Gal-8 influences primary cilium (PC) structure ARC nucleus neurons, we compared PC length in Gal-8 knockout (Gal-8 KO) mice, which lack Gal-8 expression (26, 27), and wild-type (WT) mice. We also evaluated the effect of intranasally administered Gal-8, a non-invasive route for delivering drugs and biologics into the central nervous system with limited peripheral effects (49, 50). Immunostaining of AC3 used as a specific marker for neuronal PC (14, 19) revealed larger PC in the ARC hypothalamic neurons of Gal-8 KO compared to wild-type mice, without significant differences in the percentage of ciliated neurons (Figure 1, A-C). The average PC length increased from 8.36 µm in wild-type mice to 10.64 µm in Gal-8 KO mice (Figure 1, A,B). Analysis of ciliary length distribution showed that WT mice exhibited 17.6% of cilia in the 0–4.9 µm range, 48.8% in the 5–9.9 µm range, and 33.6% exceeding 10 µm, whereas Gal-8 KO mice displayed 5.1% in the 0–4.9 µm range, 41.6% in the 5–9.9 µm range, and 53.3% over 10 µm (Figure 1C). Following intranasal Gal-8 treatment (150ng per day for 4 days), Gal-8 KO mice exhibited an average PC length comparable to that of WT mice, accompanied by a redistribution of cilia lengths to 29.6% in the 0–4.9 µm range, 35.6% in the 5–9.9 µm range, and 34.8% over 10 µm (Figure 1A-C). Thus, intranasal Gal-8 administration partially restored the ciliary length distribution in Gal-8 KO mice, particularly within the 0–4.9 µm and 5–9.9 µm ranges.

**Figure 1.**
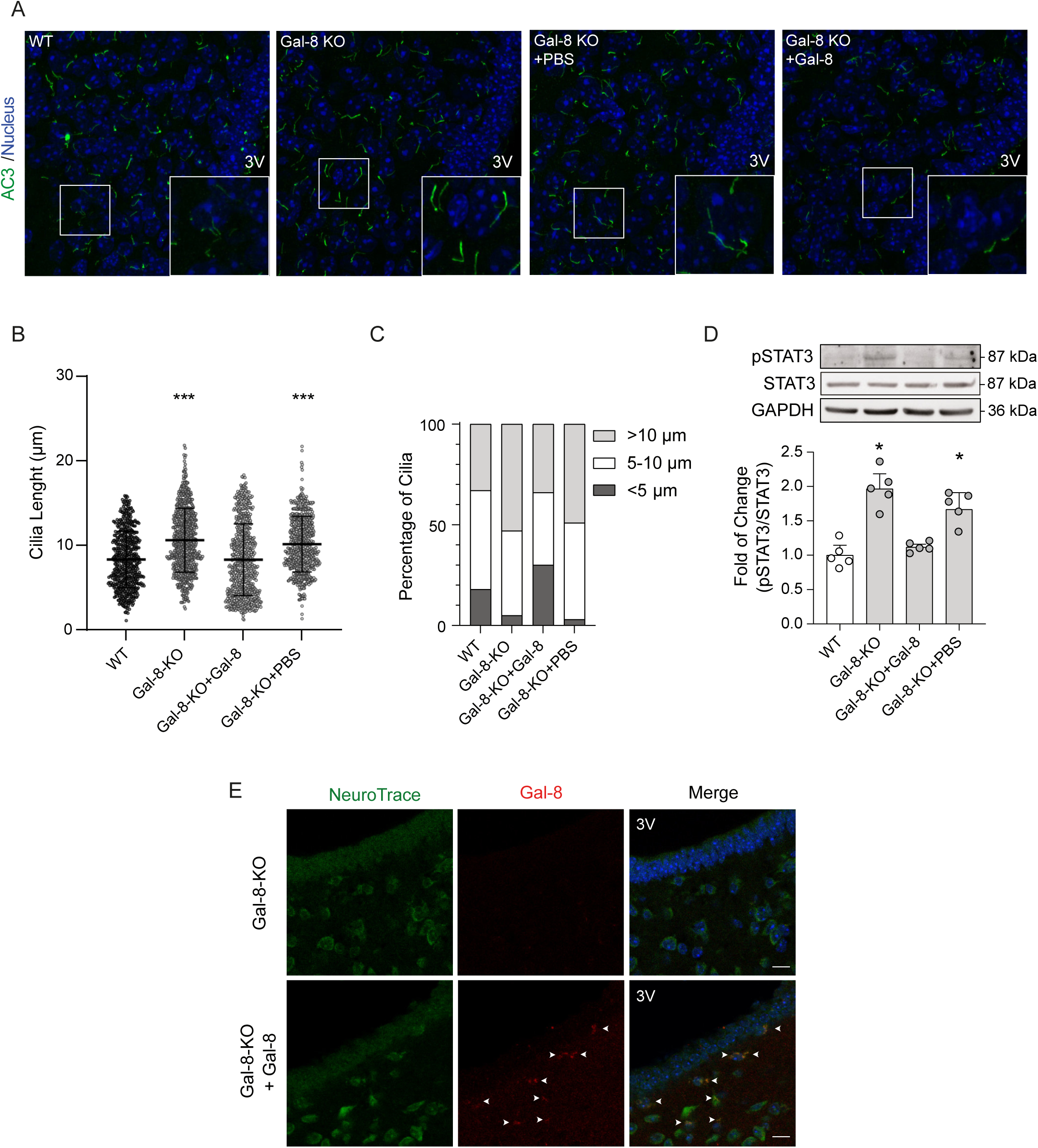
PC length, STAT3 signaling and Respiratory Exchange Ratio (RER)in Gal-8 KO mice after intranasal Gal-8 treatment. WT, Gal-8-KOmice and Gal-8-KO mice non- and treated with daily intranasal administration of PBS alone (vehicle) or Gal-8 (50 ug/ml in PBS) were analyzed for: A) Confocal images of neuronal adenylate cyclase 3 (green) immunoreactivity in the hypothalamus arcuate nucleus; B) PC length; C) PC length distribution(WT and Gal-8-KO n=7; Gal-8-KO+PBS and Gal-8-KO+Gal8,n=5; ***P<0.001; One-way Anova, SD); D) Immunoblot of STAT3 phosphorylation(***P<0.001; One-way Anova, SEM); and E) Immunohistochemistry against Gal-8 (red) and neuron staining with NeuroTrace (green) of arcuate nucleus cryosections, showing that intranasally added Gal-8 reaches the hypothalamus(n=5).

Phosphorylation of STAT3 is commonly used as a readout of leptin receptor signaling activity in the hypothalamus (7). Given that PC could be involved in leptin response (13), we next assessed hypothalamic STAT3 phosphorylation levels in Gal-8 KO and WT mice. Gal-8 KO mice exhibited a 1.8-fold increase in hypothalamic STAT3 phosphorylation compared with WT mice (Figure 1D). In contrast, intranasal administration of Gal-8 to Gal-8 KO mice reduced hypothalamic STAT3 phosphorylation to levels comparable to those observed in WT mice (Figure 1D). Immunohistochemical detection of Gal-8 in the arcuate nucleus further confirmed that intranasally administered Gal-8 reached the hypothalamus (Figure E). Together, these results indicate that endogenous Gal-8 modulates both primary cilium structure and STAT3 signaling in the hypothalamus.

### Gal-8 deficiency alters feeding behavior, locomotor activity, and body composition

Given that the leptin–melanocortin circuit regulates appetite and energy expenditure (1, 3, 51), we next compared metabolic and behavioral parameters in wild-type (WT) and Gal-8 knockout (Gal-8 KO) mice. Gal-8 KO mice exhibited lower body weight, reduced food intake, and increased locomotor activity compared with WT mice (Figure 2A–C). We then evaluated the contribution of Gal-8 to fuel selection by measuring the respiratory exchange ratio (RER) in Gal-8 knockout (Gal-8 KO) and wild-type (WT) mice. RER, defined as the ratio of CO₂ production (VCO₂) to O₂ consumption (VO₂), is commonly used to estimate relative carbohydrate versus lipid utilization (52). Gal-8 KO mice exhibited a significantly higher RER (0.95) compared with WT mice (0.87), indicating a shift toward increased carbohydrate utilization (Figure 2D). Consistent with this metabolic profile, intranasal administration of Gal-8 reduced the RER of Gal-8 KO mice to 0.88, closely resembling the WT phenotype (Figure 2D). These results indicate that Gal-8 deficiency alters whole-body substrate utilization, an effect that is reversible upon intranasal Gal-8 administration.

**Figure 2.**
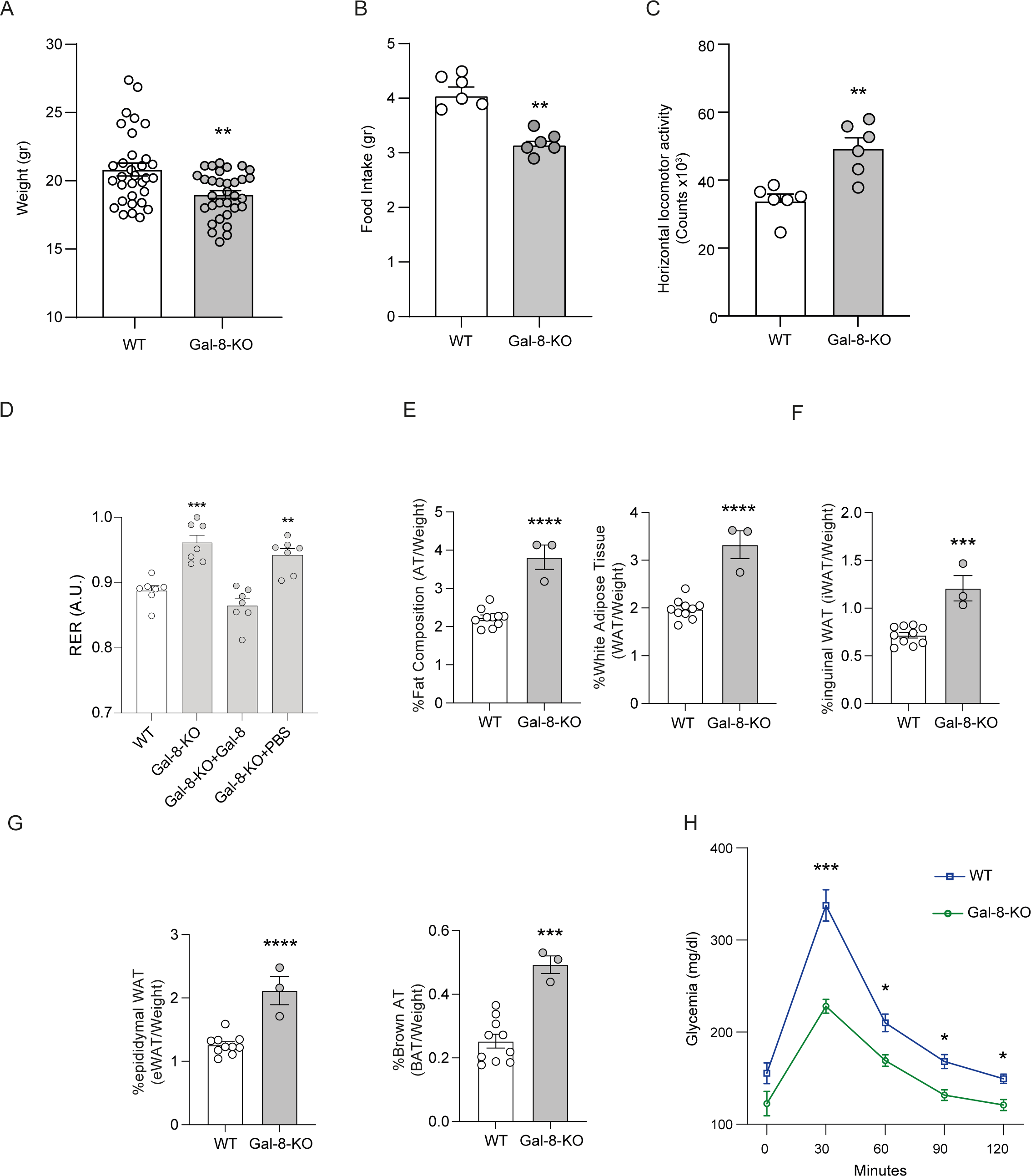
Gal-8-KO mice have metabolic alterations. Gal-8 KO mice show lower weight (A) and food intake (B), as well as an increased locomotor activity (C) compared with WT mice (n=7; P<0.01**, P<0.05*; t-test, SEM) and D) Respiratory exchange ratio (RER) of WT and Gal-8-KO mice (n=7, ***P<0.001; One-way Anova, SEM).E)Body fat composition of WT and Gal-8-KO mice. Total adipose tissue (AT) percentage and White adipose tissue (WAT). F-G) Inguinal white adipose tissue (iWAT) percentage, Epididymal white adipose (eWAT) percentage and Interscapular brown adipose tissue (BAT). WT n=10; KO n=3 biologically independent animals. H) Glucose tolerance test shows lower pick at 30 min and faster decrease of glucose levels in Gal-8 KO compared with WT mice (n=7, ***P<0.001, *P<0.05; Two-way Anova, SEM).Data are presented as mean ± SEM values and differences were analyzed with unpaired t-test, statistical significances correspond to ***P≤0.001, ****P≤0.0001.

Analysis of body composition in chow-fed animals revealed significant differences between genotypes (Figure 2E-G). Besides the reduction in the body weight, Gal-8 KO mice displayed an increased percentage of total adipose tissue, including total white adipose tissue, inguinal (subcutaneous) white adipose tissue (iWAT), epididymal (visceral) white adipose tissue (eWAT), and interscapular brown adipose tissue (BAT), relative to WT mice. These findings indicate that Gal-8 deficiency affects feeding behavior, physical activity, and adipose tissue distribution.

Since PC function in ARC neurons is implicated in the regulation of insulin signaling and glucose metabolism (1, 3, 51), we next performed glucose tolerance tests (GTTs) in Gal-8 KO and WT mice. Following intraperitoneal glucose administration, Gal-8 KO mice exhibited significantly lower blood glucose levels compared with WT mice. Glycemia was reduced by 32.4% at 30 minutes post-injection and remained lower at later time points, with decreases of 19.5%, 21.7%, and 19.1% at 60, 90, and 120 minutes, respectively (Figure 2H)

These in vivo findings support a role for endogenous Gal-8 in regulating primary cilium structure in hypothalamic neurons, with consequences for metabolic and glucose-related regulation.

### Gal-8 induces PC disassembly and resorption in hypothalamic neuron model

To investigate the cellular mechanisms regulating primary cilium (PC) length in hypothalamic neurons, we used CLU-177 cells (also referred to as mHypoA-2/12 cells), an immortalized murine hypothalamic neuronal cell line derived from adult mouse hypothalamus (44). This cell line retains functional leptin receptor signaling and responds to leptin stimulation through activation of canonical JAK2–STAT3 pathways, while expressing feeding-related neuropeptides such as NPY and AgRP (44, 53).

We first assessed the effects of Gal-8 on primary cilium (PC) presence, length, and structure in CLU-177 cells. Based on Gal-8 concentrations reported in human serum, which range from 0.18 to 5 nM in healthy individuals and from 1.34 to 15 nM in patients with proinflammatory conditions or cancer (54, 55), we tested multiple Gal-8 concentrations and identified 30 nM as sufficient to elicit clear effects on PC parameters without inducing cell proliferation (Figure S1). Treatment with Gal-8 (30 nM) for 2 h reduced the percentage of ciliated cells from approximately 80% to 60% (Figure 3A). In the remaining ciliated cells, PC length decreased from an average of 3.64 µm to 2.82 µm, accompanied by a 33.5% reduction in ciliary volume (Figure 3B–D).

**Figure 3.**
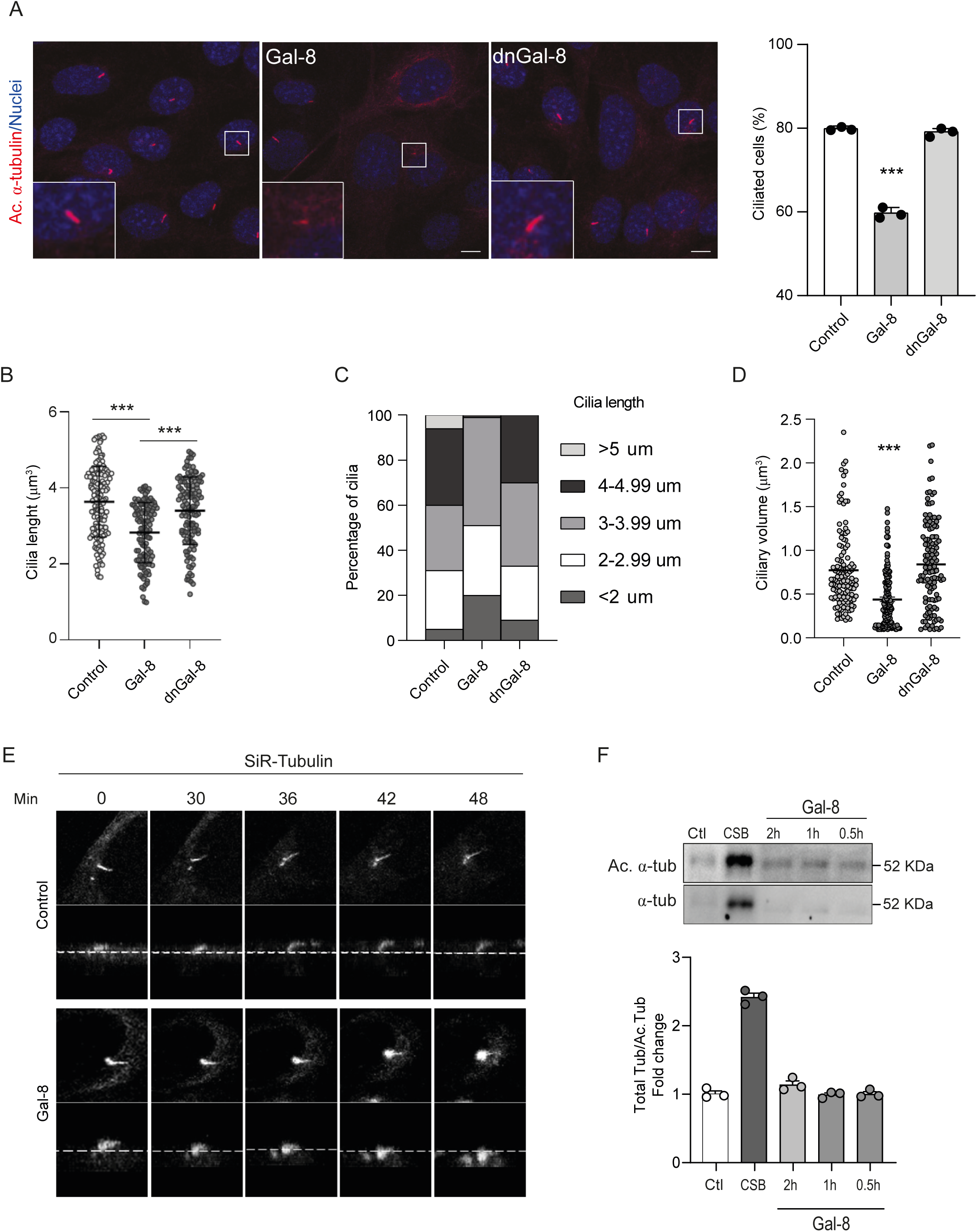
Galectin-8 induces resorption of PC in CLU-177cells. A) Indirect immunofluorescence against acetylated α-tubulin (red) showing the PC in Clu-177cells treated with 30nM of Gal-8 or denatured Gal-8 (dnGal-8) for 2 hours. Gal-8, but not dnGal-8, reduced the number of ciliated cells from 79.9% to 59.8% (P<0.001; One-way Anova, SEM); B-D) Gal-8, but not dnGal-8, treatment decreased PC length (B and C) and ciliary body volume (D). E) Live cell imaging of Clu-177cells stained with SiR-Tubulin shows PC resorption in cells treated with 30 nM Gal-8 compared with untreated cells; F) Cells stained against acetylated α-tubulin after incubation in conditioned media of cells treated with 30 nM Gal-8 or Cilia Shedding Buffer (CSB)at the indicated time points. (P<0.001; One-way Anova, SEM).

To determine the mechanism underlying Gal-8–induced PC loss, we combined live-cell imaging using the SiR-tubulin probe with detection of acetylated α-tubulin in conditioned media, a marker of ciliary shedding (17). Live-cell imaging revealed progressive PC resorption, detectable within the first 30 min of Gal-8 treatment (Figure 3E). In contrast, immunoblot analysis of conditioned media failed to detect acetylated α-tubulin following 2 h of treatment with 30 nM Gal-8 (Figure 3F). As a positive control for ciliary shedding, treatment with chloral hydrate (CH), a known inducer of PC release (17), resulted in a 2.4-fold increase in the ratio of acetylated α-tubulin to total tubulin in the conditioned media (Figure 3F).

Together, these results indicate that exposure to 30 nM Gal-8 primarily induces PC disassembly and resorption rather than direct ciliary shedding into the extracellular medium.

### Gal-8–induced primary cilium loss in hypothalamic neurons model involves activation of β1-integrin–dependent signaling

To determine whether PC loss depends on Gal-8 interactions with cell-surface glycans, we performed competition experiments in which Gal-8 was preincubated with either α-lactose or α2,3-sialyllactose (2,3SL) (28, 32, 56). α-Lactose, which competes with glycan binding to both the N- and C-terminal carbohydrate-recognition domains (CRDs) of Gal-8, completely abrogated Gal-8–induced PC loss. In contrast, 2,3SL, which selectively competes with the N-terminal CRD of Gal-8 (28, 56), partially reduced PC loss from approximately 19% to 9.7% (Figure 4A). Neither sucrose nor α2,6-sialyllactose (2,6SL) affected the ability of Gal-8 to induce PC loss (Figure 4A). Together, these results indicate that Gal-8–induced PC loss requires glycan-dependent interactions with cell-surface components.

**Figure 4.**
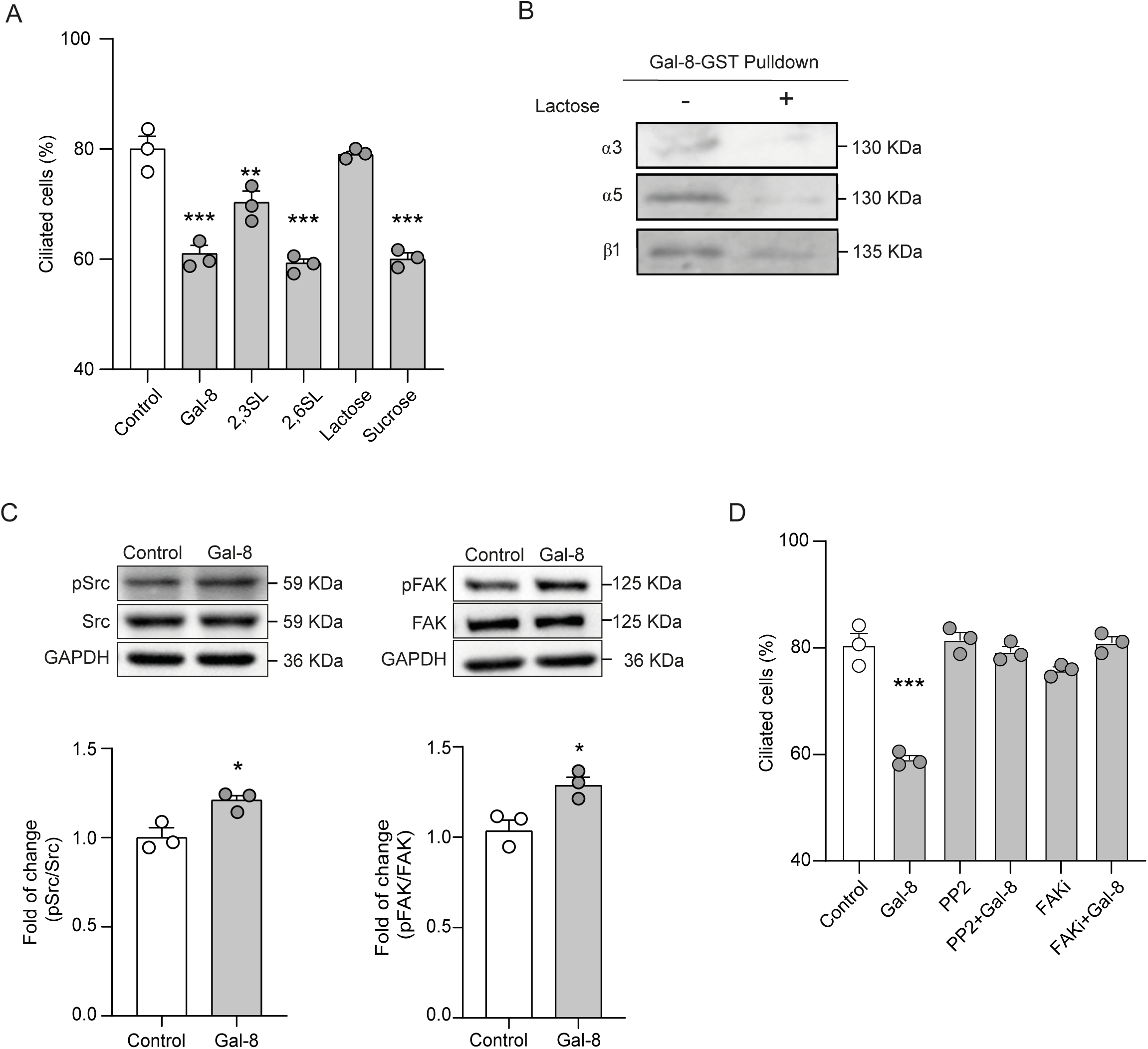
Gal-8-induced loss of PC involves interactions with cell surface glycans and b1-integrins followed by FAK/Src activation. A) Percentage of ciliated Clu-177cells treated with Gal-8 (30 nM) in the absence or presence of 10 mM α-2,3-syalillactose (2,3SL), α-2,6-syalllactose (2,6SL), or lactose. Only 2,3SL and lactose prevented Gal-8-induced PC loss. (**P<0.01 or ***P<0.001; One-way anova, SEM). B) Gal-8-GST pulldown assay shows a Gal-8 interaction with α3, α5 and β1 integrin subunit; C and D) Gal-8 (30nM, 5 min.) treatment for increased Src and FAK phosphorylation in Clu-177cells. (P<0.05; t-test, SEM). E) Src and FAK inhibitors, PP2 and FAKi, respectively, prevented PC loss induced by Gal-8 (30nM, 2h) treatment (P<0.001, One-way Anova, SEM).

Previous studies in multiple cell types have shown that Gal-8 interacts with integrins (23, 25, 30, 32) and induces activation of focal adhesion kinase (FAK), a key signaling component downstream of integrins (26, 31, 32). In addition, Src kinase, another component of integrin-associated focal adhesion complexes, has been reported to inhibit ciliogenesis and reduce primary cilium (PC) size in different cellular contexts (57, 58).

To examine whether these pathways operate in hypothalamic neurons, we performed pull-down assays and found that Gal-8 interacts with integrin α3 and integrin α5 in CLU-177 cells (Figure 4B). Immunoblot analysis further revealed that Gal-8 rapidly activated both Src and FAK, with increases of 21% and 28%, respectively, detectable within 5 min of stimulation (Figure 4C). Preincubation of cells with the Src inhibitor PP2 (20 µM) or the FAK inhibitor Y15 (10 µM) prior to Gal-8 exposure prevented Gal-8–induced PC loss (Figure 4D).

Together, these results indicate that Gal-8–mediated PC resorption in CLU-177 cells involves integrin engagement and activation of downstream FAK and Src signaling.

### Gal-8–induced β1-integrin/FAK/Src signaling promotes LTCC-mediated calcium influx, AURKA/HDAC6 activation, and primary cilium loss

Gradual PC resorption has been shown to depend on calmodulin-mediated activation of Aurora A kinase (AURKA) downstream of elevations in cytosolic calcium (38–40). Activated AURKA, in turn, phosphorylates and activates histone deacetylase 6 (HDAC6) (16, 41), which deacetylates and destabilizes α-tubulin, promoting axonemal depolymerization and PC resorption (38, 40). We therefore examined whether this pathway operates in our cell model following Gal-8 stimulation.

Ratiometric calcium imaging revealed that Gal-8 treatment induced a rapid increase in cytosolic calcium concentration, peaking within 10–15 s (Figure 5A). Chelation of extracellular calcium by addition of EGTA to the culture medium completely abolished this calcium rise and concomitantly prevented Gal-8–induced PC loss (Figure 5A-B). These results indicate that Gal-8–induced PC resorption requires calcium influx.

**Figure 5.**
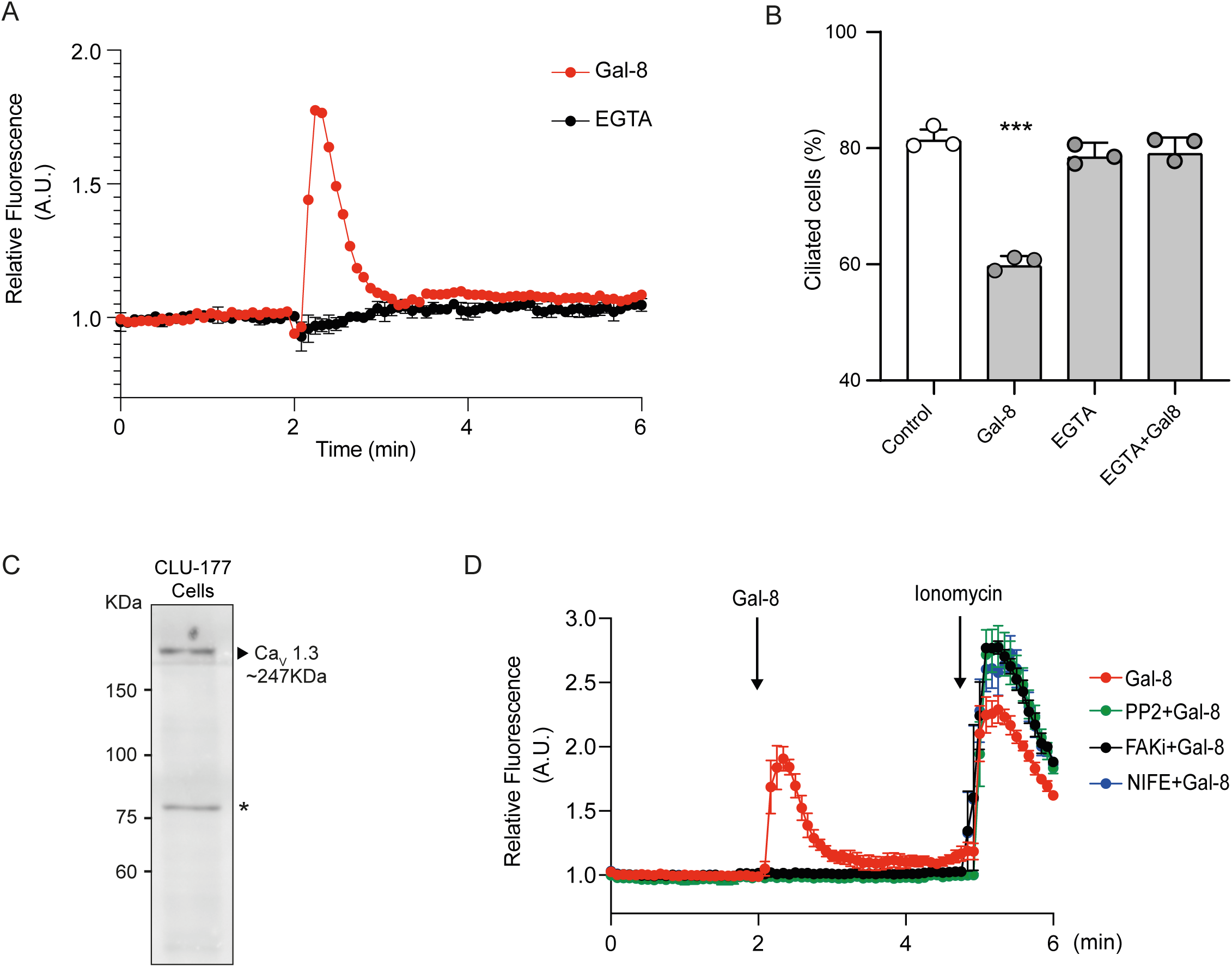
Gal-8 induces PC resorption through LTCC-associated calcium influx. A) Cytosolic calcium levels measured by Fura-Red AM imaging. Gal-8 treatment induced an increase in intracellular calcium levels, which are abrogated by EGTA, thus indicating a calcium influx; B) The PC loss elicited by Gal-8 is avoided by co-treatment with EGTA, indicating its dependency on the induced calcium influx (***P<0.001; One-way ANOVA, SEM);C) LTCC (L-type calcium channel Cav 1.3)expression. The immunoblot shows the LTCC regulatory subunit α1Din extracts of Clu-177 cells; D) Pretreatment with L-type calcium channel blocker Nifedipine (NIFE), as well as Src and FAK inhibitors (PP2 and FAKi, respectively), abrogates the calcium influx induced by Gal-8 treatment.

Activation of α5β1 integrin has been reported to induce calcium influx through opening of L-type calcium channels (LTCCs) in a Src- and FAK-dependent manner (34, 35). We detected expression of the LTCC isoform CaV1.3 in CLU-177 cells (Figure 5C). Pretreatment with the LTCC blocker nifedipine (20 µM) completely abrogated the Gal-8–induced increase in cytosolic calcium levels (Figure 5D). Inhibition of FAK and Src using Y15 (10 µM) and PP2 (20 µM), respectively, similarly prevented Gal-8–induced cytosolic calcium elevations (Figure 5E).

Consistent with activation of the calcium-dependent ciliary resorption pathway, Gal-8 treatment increased HDAC6 phosphorylation by approximately 45% (Figure 6A). Pretreatment with the AURKA inhibitor VX-680 prevented both HDAC6 phosphorylation and Gal-8–induced primary cilium loss (Figure 6A-B). Notably, blockade of calcium entry with nifedipine also inhibited Gal-8–induced HDAC6 activation (Figure 5C).

**Figure 6.**
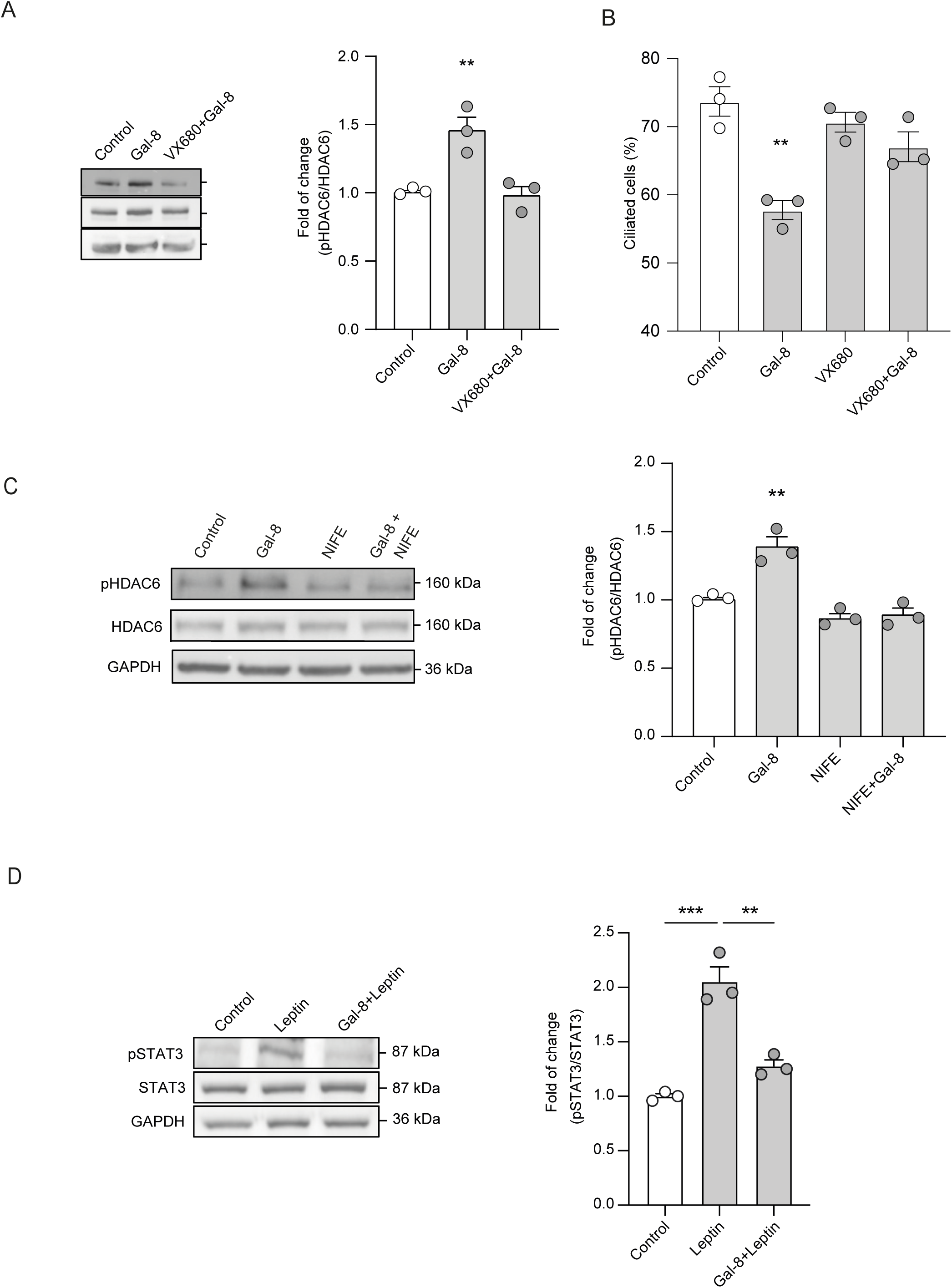
Gal-8-induced PC loss depends on calcium-mediated activation of the AurkA/HDAC6 axis. B) Immunoblot of AURKA-dependent HDAC6 phosphorylation at S22. Cells treated with Gal-8 for 2 h show an increased phosphorylation of HDAC6 at S22, which is impeded by the AurkA inhibitor VX-680, indicating that Gal-8inducesHDAC6 activation downstream AurkA activity (**P<0.01; One-way anova, SEM); B) Immunofluorescent analysis of PC presence in Clu-177 cells. The PC loss induced byGal-8 (30nM) treatment for 2 hours decreases under pretreatment with AurkA inhibitor VX-680 (P<0.01; One-way Anova, SEM); C) Calcium channel inhibitor, Nifedipine, abrogates the HDAC6 activation induced by Gal-8 (**P<0.01; One-way Anova, SEM).D) Immunoblot against phosphorylated STAT3 (pSTAT3), as a measurement of Leptin-induced signaling. Pre-treatment with Gal-8 for 2 h, which leads to PC loss as in A), decreases STAT3 activation in response to Leptin (100 nM) stimulation for 1hour. Graph shows quantification of the effect (**P<0.01, ***P<0.001; One-way Anova, SEM).

Together, these results indicate that Gal-8 induces LTCC opening downstream of β1-integrin–mediated FAK and Src activation, and that the resulting calcium influx promotes primary cilium resorption through activation of the AURKA–HDAC6 axis.

### Gal-8 attenuates leptin-induced STAT3 phosphorylation in CLU-177 cells

Alterations in primary cilium biogenesis and structural integrity in hypothalamic neurons have been shown to impair leptin signaling pathways (13, 14, 19, 20). Leptin signaling was therefore assessed by immunoblot analysis of STAT3 phosphorylation at Tyr705, a downstream event dependent on LepRb activation (59). Stimulation of CLU-177 cells with leptin (100 nM) induced an approximately 73% increase in STAT3 phosphorylation. In contrast, pretreatment with Gal-8 markedly reduced leptin-induced STAT3 phosphorylation to approximately 17% (Figure 6D).

These results indicate that Gal-8 treatment diminishes leptin/LepRb signaling in hypothalamic cells, in association with Gal-8–induced primary cilium resorption.

## Discussion

This study provides evidence of Gal-8 as a novel regulator of metabolic homeostasis. The characterization of Gal-8-KO mice phenotype together with studies in the hypothalamic cell line CLU-177/mHypoA-2/12 indicate that Gal8 is required to maintain adequate fuel use and adipose tissue content by modulating the structure and signaling properties of PC in hypothalamic neurons. Gal-8 KO mice exhibited longer PCs and increased STAT3 phosphorylation, reflecting an enhanced leptin signaling in the hypothalamus, together with alterations in hypothalamic-regulated metabolic parameters, which can be normalized by intranasal Gal-8 administration. A mechanistic approach in the CLU-177/mHypoA-2/12 cells shows that Gal-8 induces PC disassembly and resorption via LTCC-mediated calcium influx and a subsequent activation of the AurkA/HDAC6 axis, downstream of the β1-Integrin/FAK/Src signaling pathway. Gal-8 and AURKA/HDAC6 pathways may offer new therapeutic opportunities for metabolic disorders derived from hypothalamic PC dysfunctions.

We found that Gal-8-KO mice are leaner, eat less, and have higher locomotor activity than WT mice, but display increased accumulation of both white and brown adipose tissue. This suggests that the absence of Gal-8 alters energy utilization rather than caloric intake. In line with this possibility, Gal-8-KO mice have elevated RER, indicating of a shift towards the less efficient glycolytic metabolism compared with oxidative metabolism. This metabolic shift may contribute to an adipose tissue accumulation secondary to impaired fatty acid oxidation.

These findings correlate with an exacerbated PC length and function, which would promote leptin signaling towards appetite reduction and increased energy expenditure, contrasting with shortened PCs (13, 14, 20). In Gal-8-KO mice, close to 58% of the neurons in the arcuate nucleus of the hypothalamus, probably including both POMC and AgRP neurons, show longer PCs with length in the range of 10-20 μm. PCs in neurons of the arcuate nucleus contribute to insulin signaling and, thus, regulation of glucose metabolism (3, 19, 51). We found improved glucose tolerance in the GTT of Gal-8 KO mice. This property, together with the preferential glycolytic RER may be attributed to higher sensibility to leptin and insulin stimuli in hypothalamic neurons. These results suggest a role for endogenous Gal-8 in the maintenance of appropriate PC length in hypothalamic neurons, possibly by counteracting endogenous stimuli of ciliogenesis, such as leptin and insulin (14, 20).

Interestingly, blood miRNAs that control Gal-8 expression have been found to be reduced in obesity, predicting higher Gal-8 expression levels, yet in unknown tissues (60–62). PC alterations in neurons that express leptin receptor in other brain regions (3, 51, 63), or PC-dependent insulin and MC4R signaling (3, 19, 51, 64), as well as possible unknown functions of Gal-8 in peripheral tissues (23, 30), may contribute to the metabolic phenotype of Gal-8 KO mice.

Intranasal administration is a non-invasive route to deliver drugs or biologically active components into the CNS with limited peripheral spread (49, 65). This route has been widely characterized for insulin effects on cognitive dysfunctions of Alzheimer’s patients (49) and food intake in healthy individuals (66). In rats with metabolic syndrome, nasal insulin normalizes glycemia and reduces weight (67). Proteins can rapidly reach the hypothalamus after nasal administration, as shown for radiolabeled-albumin, which becomes detectable 5 minutes after administration and reaches its peak concentrations in 1 hour (50). We evaluated the effect of intranasal administration of Gal-8 in KO mice and found it to restore PC structure, STAT3-related activity, and RER close to the values observed in WT mice. These results reinforce the possibility that Gal-8 acts as a new regulator of PC-dependent metabolic balance in the brain.

The longer PC found in Gal-8-KO mice suggests that Gal-8 might negatively regulate cilia length. Our experiments in CLU177 cells show that just within 2 hours of treatment, Gal-8 reduces by ∼60% the number of cells bearing PC and decreases by ∼22.5% the length of their remaining PC. The evidence suggest resorption instead of shedding as the main mechanism of Gal-8-induced PC loss, as we found progressive PC shortening and undetectable acetylated α-tubulin as a PC component in the medium. This contrasts with the shedding process described as the prevalent PC-loss mechanism induced by serum (17). The difference very likely reflects distinct intracellular pathways affecting PC architecture. Serum stimulation likely involves PDGF and other components (38), including lysophosphatidic acid and PI3K/AKT activation (68), whereas Gal-8 mainly activates β1-integrin-mediated signaling towards FAK and Src (26, 27, 31, 32).

We show that β-lactose, currently used to block galectin interactions with glycans (26, 27, 31, 32), abrogates Gal-8-induced PC loss, thus demonstrating its dependency on Gal-8 interactions with cell surface glycans. Gal-8’s N-terminal carbohydrate recognition domain (CRD) prefers terminal α-2,3-sialyllactose, whereas its C-terminal CRD shares with other galectins the general preference for N-acetylactosamines (28, 56). We also show that α-2,3 but not α-2,6 sialyllactose prevents Gal-8-induced PC loss, indicating a selective requirement of glycan interactions with Gal-8’s N-terminal CRD.

Integrins are the main counter-receptors of Gal-8 (23, 25, 30), and their activation, followed by downstream FAK/Src kinase signaling, regulates diverse cellular processes (23, 25, 27, 30–32). In line with previously described interactions of Gal-8 with cell surface glycoproteins in other cellular systems (26, 27, 31, 32), including hippocampal neurons (27), our pull-down experiments show that Gal-8 preferentially binds α3- and α5- but not α4- β1-integrins in CLU177 cells. Accordingly, we found that Gal-8 treatment activates FAK and Src and also that their inhibitors prevent the PC loss. These results implicate the β1integrin/FAK/Src pathway in the PC resorption process.

Previous studies describe that activation of α5β1 integrins with ligands such as RGD peptides, fibronectin, or anti-a5 antibodies induces calcium influx via LTCC downstream of FAK and Src in basal forebrain neurons and vascular smooth muscle cells, two types of excitable cells (34, 35). This effect involves an α5β1 interaction with LTCC (33). Also, overexpression of α3β1-integrin, as found in tumor-associated endothelial cells in glioblastoma, promotes calcium influx, though the mechanism has not yet been elucidated (37). Gal-8 itself induces calcium influx in platelets via an unknown mechanism (69). Here, we show that Gal-8 triggers calcium increments in CLU177 cells, mitigated by EGTA, implying calcium influx. This calcium influx is abrogated by the LTCC inhibitor Nifedipine, as well as by FAK or Src inhibitors, indicating that Gal-8 triggers the described α5β1/FAK/Src/LTCC pathway resulting in calcium influx in CLU177 (34, 35). As expected, this calcium influx activates AURKA and HDAC6, and their inhibition, as well as all conditions that prevent calcium influx, impedes PC loss induced by Gal-8 in CLU177 cells. AURKA is known to phosphorylate the deacetylase HDAC6, enhancing its deacetylase activity, which deacetylates the polymerized α-tubulin of the PC axoneme and promotes its disassembly without affecting non-ciliary acetylated microtubules in the cytoplasm (38). HDAC6 also deacetylates cortactin, which leads to actin polymerization, additionally contributing to PC disassembly (70). Altogether, these data indicate that Gal-8, acting through a selective β1-integrin/SRC/FAK/LTCC pathway, triggers a well-known mechanism of PC resorption involving AURKA/HDAC6 activation by cytosolic calcium increase. Our results are the first to demonstrate that Gal-8 activates this mechanism of PC regulation.

Leptin controls feeding, maintains energy homeostasis, and regulates glucose metabolism by activating its receptor LepRB, which is highly expressed by antagonistic POMC and AgRP neurons in the arcuate nucleus of the hypothalamus, as well as by neurons of other brain regions involved in neuroendocrine control (51, 63). Leptin-activated LepRB triggers downstream signaling pathways that include JAK2-STAT3 (71). Previous work identified LepRB at the base of the PC and suggests that functional PCs are necessary for appropriate leptin signaling in the hypothalamus (13, 14). Our results show that Gal-8 pretreatment decreases leptin signaling towards STAT3. Therefore, Gal-8 can potentially regulate PC-dependent leptin functions affecting food intake and energy balance (13).

Previous studies using Gal-8-KO mice and experimental autoimmune encephalomyelitis, a model of multiple sclerosis, suggested a role of Gal-8 as immunosuppressor protecting against damaging neuro-inflammatory conditions in the brain (26). A neuroprotective role of Gal-8 has also been proposed from experiments in neuronal primary cultures and stereotaxic injection of a damaging agent such as H_2_O_2_ in hippocampus (27). We have described the presence of circulating anti-Gal-8 autoantibodies in systemic lupus erythematosus (SLE), rheumatoid arthritis, during sepsis (72) and in multiple sclerosis (MS) (26), which can be function-blocking (26, 31, 73, 74), and able to neutralize the immunosuppressive and neuroprotective roles of Gal-8 (26, 27). Their potential pathogenic roles have so far been associated with lymphopenia in SLE (72) and with worse prognosis in patients with the relapsing-remitting form of MS (26). It is also possible that circulating anti-Gal-8 autoantibodies reach the hypothalamus by diffusing from the fenestrated capillaries of the nearby medial eminence (75) and block the function of Gal-8 on the neuronal PC, leading to metabolic alterations.

Glycan structural changes that affect the interactions of galectins with cells can occur in physiological or pathological conditions (76, 77). The so-called “glycan code” is dynamically translated by lectins, including galectins, in different organs and systems (25, 76–78), associated with metabolic alterations, including obesity and alteration in glucose homeostasis (76). N-acetylglucosamine (GlcNAc), which promotes N-glycan branching and affinity changes for galectins (76, 79), have been found to correlate with insulin resistance and weight gain in mice when dietary supplement or derived from the gut microbiota (79, 80). On the other hand, secreted neuraminidases release sialic acid from cell surface glycans and thus decrease their interactions with Gal-8 (81–84). Peripheral organs such as the liver show increased sialylated glycans in obesity (85, 86). Mice receiving a high-fat diet display reduced expression of neuraminidase Neu1 associated with insulin resistance and hyperglycemia (87). Interplays between N-Glycan branching and sialylating/desialylating conditions affecting Gal-8-dependent PC functions in the hypothalamus, need to be investigated in the future.

Non-mitotic AURKA roles have been increasingly explored since its first identified involvement in PC disassembly during G/G1 cell cycle phase (41). However, the only function of AURKA described in post-mitotic neurons is related to neurite extension, which involves PKC–AURKA–NDEL1 signaling axis (41, 88). The crucial role of the PC in transducing metabolic signals from leptin, insulin and MC4R (13, 14, 19, 64), suggests that AURKA, together with its variety of protein interactors, regulators, and effectors (16, 41, 89), may be important players in the regulation of the neuroendocrine system, with potential therapeutic applications. Genetic human disorders and preclinical models of some ciliopathies may be directly connected to the regulation of AURKA (40–42). For instance, INPP5E, a protein defective in MOMR ciliopathic syndrome that includes obesity, seems to form a positive feedback loop of mutual activation with AURKA, suppressing ciliogenesis (41, 42). Ciliogenic promoting actions of leptin and insulin (14, 20), as well as the recently described shortening and loss of the PC in hypothalamic neurons exposed to palmitic acid or high-fat diets leading to insulin resistance (19), may involve known regulators of the AURKA/HDAC6 axis.

New translational opportunities may arise from our results. For instance, specific AURKA inhibitors used in clinical tests against cancer (41), and L-type calcium channel inhibitors, such as Nifedipine used in hypertension (90), might be repurposed to promote hypothalamic ciliogenesis associated with the control of appetite and metabolic homeostasis. A similar approach may stimulate searching for specific Gal-8 inhibitors (91). Exploration of intertwined AURKA and Gal-8 pathways could lead to innovative therapies for metabolic disorders, emphasizing the significance of these proteins in maintaining metabolic homeostasis and offering new avenues for clinical intervention.

## Methods

### Animals

Male mice, 3-4 months old, were housed at the Faculty of Biological Sciences of the Pontificia Universidad Católica de Chile, and maintained under conditions of strict confinement, including automatic control of temperature (21°C) and photoperiod (12 h light / 12 h dark), with water and food ad libitum. All protocols were approved by the Institutional Ethics Committee of Care and Use of Animals in Research of Universidad San Sebastián (protocol number 21-2021-20). Lgals8/Lac-Z knock-in (here Gal-8-KO) mice have Lgals8 gene (18,427 bp) replaced with LacZ lox-Ub1-EM7-Neo-lox Cassette containing the LacZ gene that encodes b-galactosidase (26, 27). Animals were randomly assigned to control or test groups for each experiment and maintained in the same environment to avoid bias. Each animal was considered an individual experimental unit. Body weight was measured using a mouse scale (Accuris Instruments, Model W3300-500). food intake and locomotor activity were measured for 12 h in the dark cycle using a Metabolic Cage (Ugo Basile S.R.L. Model 41800-010).

### Regents and materials

The following reagents were purchased from Sigma: Thrombin from human plasma (T1063), bacterial protease inhibitor cocktail (P8465), Src inhibitor PP2 (P0042), FAK inhibitor Y15 (SML0837), and protease and phosphatase inhibitor tablets (A32959, Pierce). Reagents from Santa Cruz Biotech included: Nifedipine (SC-3589), and antibodies against α-tubulin (B-5-1-2), Cav a1D (E3), a-3 (A-3), a-5 (H-104) and b1 (4B7R) integrins, FAK (H-1), pFAK (2D11), Src (B-12) and pSrc (9A6). Cell Signaling Technology provided antibodies against acetylated-α-tubulin K40 (D2063), HDAC6 (D21B10), STAT3 (D3Z26) and pSTAT3 (M9C6). Anti-pHDAC6 (Ab61058) was purchased in Abcam. Secondary antibodies conjugated with horseradish peroxidase or AlexaFluor were obtained from Rockland and Molecular probes, respectively. Glutathione-sepharose 4B (17075601) (Cytiva), isopropyl-1-thiogalactopyranoside (IPTG) (Invitrogen) SiR-Tubulin (CY-SC002;Cytoskeleton), Fura-Red AM (Cat # F3020;Invitrogen)

### Cell culture

Clu-177 cells were maintained with high glucose Dulbecco’s Modified Eagle’s medium (DMEM) supplemented with 10% Fetal Bovine Serum, 100 ug/mL penicillin, and 0,1 mg/mL of streptomycin (P/S) in a cell incubator with a constant flow of 95% O2 and 5% CO2 at 37 °C. Cells were tested for mycoplasma contamination every two weeks. Eighteen hours prior to the experiments, cells at 70% confluency were washed with PBS 1x and maintained with DMEM-P/S media without FBS to induce ciliogenesis (17).

### Recombinant Gal-8 production

Human recombinant Galectin-8 (Gal-8) was produced as previously described (27). Briefly, bacteria transformed with pGEX-4T-3 (Pharmacia Biotech) plasmid bearing Gal-8-GST or GST were grown in a pre-inoculum in LB supplemented with ampicillin (100 µg/ml) (LB/Amp) at 37°C and 200 RPM of agitation. Gal-8-GST production was induced by IPTG (0.1 mM). Bacterial pellets were lysed for total protein extraction, and Gal-8-GST was purified via a Glutathione-Sepharose column. Gal-8 was released from GST-Gal-8 linked to glutathione-Sepharose by thrombin treatment (10 U/mg of fusion protein) for 4 h at room temperature.

### Immunofluorescence

Clu-177 cells were fixed with ice-cold paraformaldehyde (PFA) and 4% sucrose for 10 min and incubated with blocking buffer (TBS/Saponin 0.05%/BSA 1%) for 30 minutes at RT. Cells were then incubated overnight with primary antibody diluted in blocking buffer at 4°C, followed by secondary antibody conjugated with AlexaFluor and Hoechst for 1 h at 37°C and finally mounted with Fluoromount. Arcuate nucleus slices, prepared as previously described (19), were incubated overnight at 4°C with the primary antibody against adenylate cyclase 3 (AC3) or Gal-8. Slices were then incubated with secondary antibodies for 1 h at room temperature and mounted with Fluoromount. Slides stained with NeuroTrace probe were dyed according to the manufacturer’s instructions.

### Analysis of PC morphology

The percentage of cells with primary cilia (PC) was evaluated by counting the number of ciliated cells versus their total number in at least 100–150 cells per sample from three different experiments. PC images were captured by z-stacking on a Leica SP8 or Zeiss AiryScan confocal microscope. PC morphology was analyzed using the FIJI plugin CiliaQ, as described by Hansen et al (45).

### Western Blots

Cells exposed to the different treatments were lysed with RIPA lysis buffer (Tris 50 mM, NaCl 150 mM, EDTA 5 mM, NP40 1%, Sodium Deoxycholate 0.5%, SDS 0.1%, pH 8.0) for 30 minutes at 4°C, supplemented with protease and phosphatase inhibitors. Samples were resolved in an SDS-PAGE gel and transferred onto a PVDF membrane using the Bio-Rad Trans-Blot Turbo system. Membranes were blocked, and primary antibodies were incubated overnight at 4°C in blocking buffer. Secondary antibodies were incubated for 1 h in blocking buffer at 37°C. Images were acquired using the Syngene G:Box detection system, and densitometric band analysis was performed using FIJI software.

### Pull-down assay

Protein extracts were preincubated with GST-Glutation Sepharose column for 1 h at 4°C, then incubated for 2 h at 4°C with Gal-8-GST-Glutation-Sepharose. The bound protein was eluted and analyzed by immunoblot, as previously described (26).

### Conditioned media analysis

Clu-177 cells were treated with Gal-8, vehicle or Shedding Cilia Buffer (46) (SCB) (112 mM NaCl, 3.4 mM KCl, 10 mM CaCl2, 2.4 mM NaHCO_3_, 2 mM HEPES, pH 7.0). The conditioned media was centrifuged at 1,000g x g for 10 min to remove cell debris, and the supernatant was incubated with 20% trichloroacetic acid (TCA) overnight at 4°C with agitation. Samples were centrifuged at maximum speed for 10 min and prepared for western-blot analysis.

### PC live cell imaging

Clu-177 cells seeded in live cell imaging plates were incubated with 100 nM SiR-Tubulin overnight, washed twice with 1mL of DMEM-Hepes, and live cell imaging was performed with a Leica SP8 confocal microscope, using a 63x oil immersion objective and time resolution of 180 seconds per frame.

### Ratiometric calcium imaging

Clu-177 cells were incubated with 5 uM Fura-red AM for 30 minutes. Pre-treatments with blockers or inhibitors were performed post incubation with Fura-red AM. Live cell imaging was performed with a Leica SP8 confocal microscope with a 63x oil immersion objective and a time resolution of 5 seconds per frame.

### Locomotor activity

Mice were placed in a metabolic cage (UGO BASILE, model 41801) for seven hours, and then their locomotor activity was measured during the night phase for 12 hours, with food and water *ad libidum*.

### Respiratory exchange ratio

CO_2_ production and O_2_ consumption were measured using the iWorx GA200 measurement system according to the manufacturer’s instructions. Mice were placed in the metabolic cage at a maximum concentration of 0.24% CO_2_. Measurements were performed until a CO_2_ concentration of 3% was achieved. Data were analyzed with the LabScribe 2.0 software according to the manufacturer’s instructions.

### Glucose tolerance test

The mice’s basal glycemia was measured after 6 hours of fasting. A glucose solution (2 g per kg of body weight) was injected subcutaneously. Glycemia measurements were performed every 30 minutes post-injection for 2 hours. During the measurement period, the mice were deprived of food to avoid interference with the measurement (47).

### Body fat composition

To evaluate body fat composition of WT and Gal-8-KO mice, after euthanasia, interscapular brown adipose tissue (BAT), inguinal (subcutaneous) white adipose tissue (iWAT) and epidydimal (visceral) white adipose tissue (eWAT) was extracted and weighed.

### Intranasal Gal-8 administration

Intranasal administration of Gal-8 was performed in 3-month-old Gal-8-KO mice partially anesthetized with isoflurane as previously described (48). In brief, in a prone position, 3 uL of Galectin-8 (50 mg/ml) or PBS was administered to the treated or control group, respectively, in each nostril. The procedure was repeated once a day for 4 days.

### Statistical analysis

Analyses were conducted with GraphPad Prism 8.0 software. Student’s t-test was used for two-group comparison, and one-way or two-way analysis of variance (ANOVA) with Tukey’s multiple comparisons test was used for comparisons of more than two groups. P-values equal to or minor than 0.05 (P≤.05) were considered statistically significant. The confidence interval range was 95% in all tests. The p-value graphic style is indicated according to APA format.

## AUTHOR’S CONTRIBUTION

**C.H.:** Conceptualization, Data curation, Formal analysis, Validation, Visualization, Investigation, Methodology, Writing**. M.H; D.P.; A.A.; F.P.; A.V.; D.C. C.J., S.E.:** Investigation, Methodology, Data curation. **E.M, B.K and C.T.:** Conceptualization, Resources, Data curation, Writing**. A.S.:** Resources, Conceptualization, Writing. **A.G.:** Founding Acquisition, Resources, Conceptualization, Data curation, Supervision, Validation, Writing.

